# What we can find in what’s left behind: DNA metabarcoding of semi-aquatic insect exuviae

**DOI:** 10.1101/2024.09.27.615410

**Authors:** SL Dworatzek, M Tozer, K Austin, A Vermey, D Steinke

**Affiliations:** Department of Integrative Biology, University of Guelph, Guelph, Ontario, Canada N1G 2W1; Centre for Biodiversity Genomics, University of Guelph, Guelph, Ontario, Canada N1G 2W1

**Keywords:** Ephemeroptera, Plecoptera, Trichoptera, Odonata, Chironomidae, Insect emergence, Non-invasive research, Wetlands, Freshwater, Benthic Invertebrates

## Abstract

Semi-aquatic insects contribute to critical ecological functioning of freshwater habitats in both the aquatic and terrestrial phase of their life. Traditional methods of studying their emergence rely on the capture of insects as they emerge, and morphological identification with taxonomic keys. This is not only time consuming but can have large impacts on the study population, obstacles that can be removed by the use of DNA.

This study investigated the potential of using exuviae collected from the water surface as DNA source. Both emergence trap samples and insect exuviae were collected from a constructed wetland and a small creek in southern Ontario. Metabarcoding provided a total of 40 samples with 254 distinct taxa from emergence traps (26 samples), and 135 from exuviae (14 samples). There were many similarities between both two sample types, especially for the rich chironomid diversity. Nonetheless, the exuviae samples were able to identify more orders containing semi-aquatic insects. Furthermore, they showed a higher level of diversity within these orders. This higher level of diversity seen in exuviae samples may be due to limitations of emergence traps, such as they only account for a small defined surface area. In contrast, exuviae are representing a much larger area and are free floating, thus collection of emerging taxa is not limited to the emerging site.

We were able to show that identification of emerging aquatic insects through metabarcoding of exuviae is a useful method for the study of insect emergence.

## Introduction

About 2% of all known insects exhibit aquatic life stages (Leveque et al., 2005). Most of them spent much of their life under water, surviving as adults only for a short period, used primarily for reproduction. When such semi-aquatic insects emerge from the aquatic environment, they undergo ecdysis, the shedding of the larval exoskeleton and formation of the final adult stage (Cheong et al., 2015). Some notable insect orders in which all or many species are semi-aquatic include Ephemeroptera, Plecoptera, Trichoptera, Odonata, Megaloptera, and Diptera (Barber-James et al., 2008; Cover & Resh, 2008; Ferrington, 2008; Fochetti & Tierno de Figueroa, 2008; Kalkman et al., 2008; Lancaster & Downes, 2013; Moor & Ivanov, 2008). Despite representing just a small subset of all insects these orders make up a large proportion of the diversity of the freshwater benthos (Balian et al., 2008). Other highly diverse orders such as Hemiptera, Lepidoptera, Coleoptera, and Hymenoptera are primarily terrestrial but still contain some semi-aquatic and fully aquatic species (Bennett, 2008; Mey & Speidel, 2008; Polhemus & Polhemus, 2008).

Many semi-aquatic insects provide critical ecosystem services. As adults, they transfer mass and energy from aquatic to terrestrial environments (Hunter & Price, 1992; Kagata & Ohgushi, 2006). It has been shown that many terrestrial vertebrate species rely on insect emergence for a large portion of their diet (Murakami & Nakano, 2002; Sabo & Power, 2002; Salvarina et al., 2018). Some species of Diptera and Ephemeroptera are considered ecosystem engineers, as they are breaking down organic matter, and mobilizing nutrients for other organisms (Adler & Courtney, 2019; Jacobus et al., 2019).

As freshwater habitats have been identified as among the most threatened ecosystems worldwide (Albert et al., 2021; Vörösmarty et al., 2010), studies of semi-aquatic insects and their response to anthropogenic change have become more important especially in the context of better understanding the factors contributing to the health of freshwater ecosystems. Benthic invertebrates are highly sensitive to changes in water quality and therefore are often used as indicator species (Bonada et al., 2006). Many species have specific limits for various environmental variables, thus their presence or absence can be indicative of the status of environmental conditions (Bustos-Baez & Frid, 2003.; Kubosova et al., 2010). Consequently, semi-aquatic insects have been used as bioindicators, predominantly to assess water quality; however, research is now shifting to the possibility of using them to measure the impact of climate change (Bunn et al., 2010; Bush et al., 2013; Khamis et al., 2014).

While most monitoring programs will focus on the collection of aquatic live stages using various trap types, the study of insect emergence typically involves standard pyramidal traps, which collect adult insects as they emerge from the water column (Mundie, 1956). These traps cover a portion of the water’s surface, catching insects as they go through ecdysis. Although this method is effective, it has its drawbacks due to its impact on insect populations and the limited surface area it covers. There is an ongoing movement to transition to non-invasive research methods, which have lower impacts on study populations (Beja-Pereira et al., 2009; Zemanova, 2020). A non-invasive alternative for collecting samples to study emergence, involves the gathering of exuviae from the surface, but to date this method has only been used for specific taxonomic groups (Arpellino et al., 2022; Hadjoudj et al., 2014) where exuviae retain enough morphological traits to allow for proper identification. In fact, most studies utilizing either emergence traps or exuviae collection relied on species identification with traditional taxonomic keys. This can be overly time consuming and given the limited number of characteristic traits of exuviae not very reliable. An alternative to this would be DNA barcoding (Hebert et al., 2003) and its extension for community samples, DNA metabarcoding (Porter & Hajibabaei, 2018). Especially the latter has been successfully applied to bulk arthropod samples, even when only trace amounts of DNA were available (Elbrecht et al., 2017; Ritter et al., 2019; Ruppert et al., 2019; Elbrecht et al., 2021; Steinke et al., 2022).

The use of insect exuviae for DNA analysis is not new and has been used for a few terrestrial groups such as cicadas, butterflies, beetles, and tarantulas (Feinstein, 2004; Kranzfelder et al., 2016, 2017; Lefort et al., 2012; Nguyen et al., 2017; Petersen et al., 2007). However, only a few studies focused on speciose semi-aquatic insect groups such as chironomids (Krosch & Cranston, 2012). Another more recent study employed DNA barcoding of Odonatan exuviae to monitor their emergence (Sittenthaler et al., 2023). Despite the relative effectiveness of this approach, they were also limitations such as DNA degradation due to environmental exposure and the presence of PCR inhibitors (Nguyen et al., 2017; Sittenthaler et al., 2023; Watts et al., 2005). While these studies demonstrated the possibility of using insect exuviae for DNA-based species identification all studies focused on a single taxon. Thus far, there has been no comprehensive study of insect emergence including exuviae of multiple taxa.

Consequently, our study uses insect exuviae collected from the water surface of a pond to determine if they contain sufficient residual (environmental) DNA to identify past and present emerging taxa. We also collected samples using a standard emergence trap. Both sample types were collected at the same sampling site, over the same time period, using identical DNA extraction, amplification and sequencing protocols allowing for a direct comparison of the two sampling methods. and how the use of DNA metabarcoding of insect exuviae might impact the study of aquatic insect emergence.

## Methods

### Sample Collection

Weekly samples were collected from a site in Kitchener-Waterloo, Ontario, Canada (43.4921828, -80.6157117) between May 29th and August 21^st^, 2023. Collection was done at a creek, and a manmade pond, which the creek fed into. For emergence trap samples, four traps were deployed, three on the pond and one on the creek. Soil emergence traps were modified to be buoyant and anchored in place (Figure 1) (Malison et al., 2010). Exuviae were collected weekly from each surface using a sieve, and then transferred to ethanol using sterile forceps. Both sample types were stored in 95% ethanol before DNA extraction.

**Figure 1.**
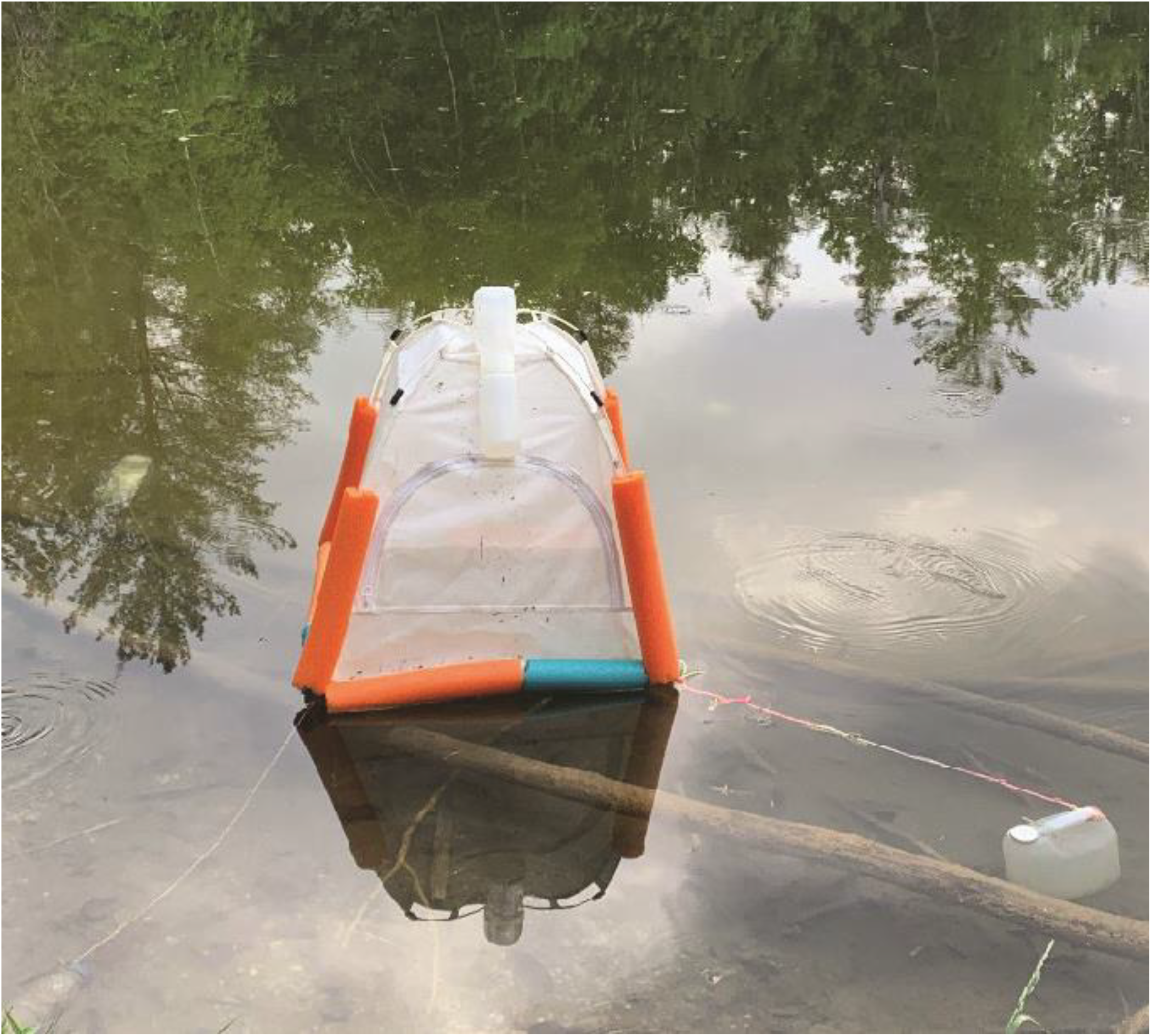
Emergence trap at study site.

### Lysis and Extraction

For both exuviae and emergence trap samples the ethanol was drained, and samples were dried in an incubator for a minimum of four hours. Less time was required to dry exuviae due to the lower biomass of these samples. Once dry, samples were ground into a fine powder using either the IKA© tube mill (IKA, Breisgau, Germany) at 25,000 rpm for 3 min or by vortex with steal beads (diameter 5mm). Lysis was done using approximately 25 mg of the powder, and 200 μl of a solution containing 90% insect lysis buffer (7000 mM GuSCN, 30 mM EDTA pH=8.0, 5% Tween-20, 30 mM Tris-HCl pH=8.0, 0.5% Triton X-100) and 10% Proteinase K (Qiagen, 20 mg/ml). Samples were vortexed vigorously and incubated at 58 °C overnight, on a shaker, before DNA was extracted using the Qiagen DNeasy Blood & Tissue Kit following manufacturer instructions.

### Amplification, library preparation, and sequencing

Amplification of the cytochrome oxidase subunit 1 (COI) gene was done using a two-step PCR protocol with the forward primer BF3 (5’-CCHGAYATRGCHTTYCCHCG-3’) and reverse primer BR2 (5’-TCDGGRTGNCCRAARAAYCA-3’) (Elbrecht & Leese, 2017; Elbrecht & Steinke, 2019). The first PCR was completed with a total volume of 25 μl, using 5 μl of the extract DNA, Multiplex PCR Master Mix Plus (Qiagen), 0.5 mM of each primer, and molecular grade water. The second PCR was conducted using fusion primers with sample specific inline tags and used 5 μl the PCR1 product (Elbrecht & Steinke, 2019). Both PCR cycles had the same conditions (25 cycles with denature 95°C 30 seconds, annealing at 50°C 45 seconds, and extension at 72°C for 50 seconds, with an additional 5 mins at 95°C at the start and 72°C at the end). The size of the amplification product was monitored after PCR2 with a 1% agarose gel and standard 1kb DNA ladder.

The total volume of PCR product was normalized using the Invitrogen SequalPrep™ Normalization plate kit (Thermo Fisher Scientific, MA, USA, Harris et al. 2010). Following this, 5 μl of each sample were pooled together in a 5 ml reaction tube, vortexed, and distributed equally into 1.5 ml microcentrifuge tubes for solid-phase reversible immobilization (SPRI)-based size selection (Beckman Coulter). The left side size selection procedure was used with a sample-to-volume ratio of 0.7 and was eluted into 20 μl of molecular grade water. The elutes were pooled into a 1.5 μl reaction tube and monitored on a 1% agarose gel before the DNA concentration was measured for five replicates with the Qubit dsDNA HS Assay Kit. The pooled sample was submitted for sequencing on an Illumina MiSeq with the 600 cycle Reagent Kit v3 and 5% PhiX spike in at the University of Guelph Advanced Analysis Centre.

### Data analysis

Sequences were demultiplexed and subjected to further filtering and analysis as outlined in Buchner et al., (2022): after processing the raw reads using the APSCALE pipeline (Buchner et al., 2022), taxonomy was assigned to Operational Taxonomic Units (OTUs) using a curated Canadian Reference Library (Pentinsaari et al, in prep). All analyses and visualizations were made using R Version 4.2.2 (R Core Team 2022) with the package “ggplot2” (Wickman 2016).

## Results

In total, 40 emergence trap samples and 26 exuviae samples were collected, of these, 28 emergence trap, and 14 exuviae samples were successfully sequenced. Damage to two of the traps limited the creek trap to the first five weeks of sampling, and one of the wetland traps to the first six weeks of sampling. The quantity of DNA extracted from emergence trap samples (Mean concentration: 21.5 ng/μl) was significantly (Chi-squared = 24.883, p-value = 6.092e-07) higher than exoskeleton samples (Mean concentration: 5.2 ng/μl). Illumina sequencing produced a total of 21,340,833 reads. Raw data are available on NCBI SRA under the accession number XXXXX. After filtering emergence trap samples, we retrieved 894,980 total reads (average number of reads per sample 31,964), and for the exuviae 75,713 total reads (average number of reads per sample 5,408). We found 254 distinct ASVs in emergence traps and 135 distinct ASVs in the exuviae samples. After correcting for the number of samples, the ratio of species per sample in emergence traps is approximately nine per sample, while for exuviae it is approximately ten per sample.

Most matches belonged to Insecta (98% emergence samples, 97% exuviae sampling – Figure 2). In emergence trap samples the order Diptera makes up 74% of insects and 72% of all detections (Figure 3) and the majority of reads obtained across all samples (Figure 4). Diptera were seen at a lower rate in exuviae samples, but are still the most prevalent, making up 60% of insects and 53% of all detections. In both sample types, most Diptera are members of the family Chironomidae with 60% of the total detections, as well as 83% of all dipteran detections in emergence trap samples (Figure 2A). In exuviae samples they make up 40% of all detections and 76% of all Dipteran detections (Figure 2B). In both sample types three Chironomidae subfamilies were detected (Chironominae, Orthocladiinae, and Tanypodinae) with the former being the most prevalent (Figure 3).

**Figure 2.**
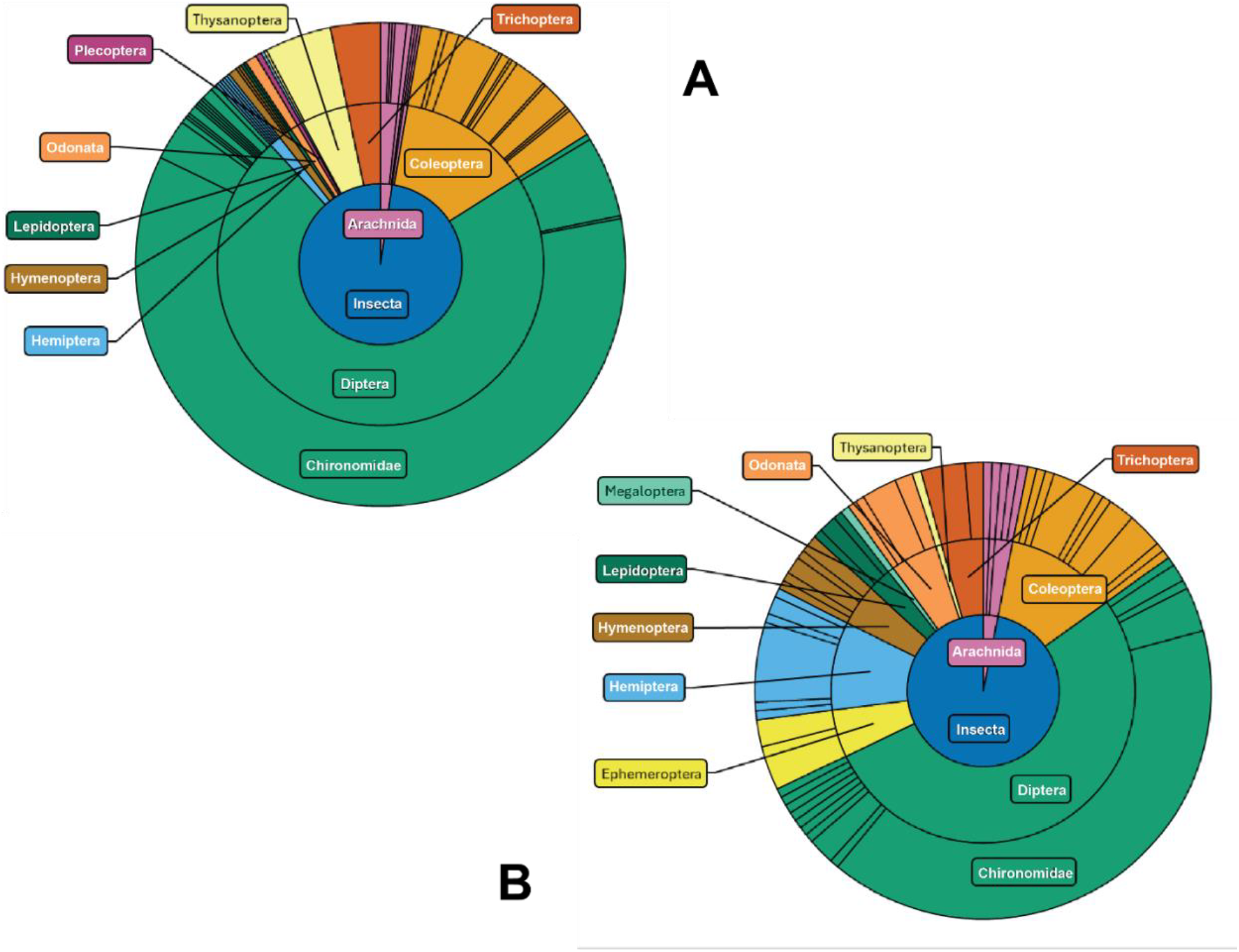
Taxonomic diversity based on the number of detections of A) the exuviae samples and B) the emergence trap samples. Important taxonomic groups are labeled with corresponding colours.

**Figure 3.**
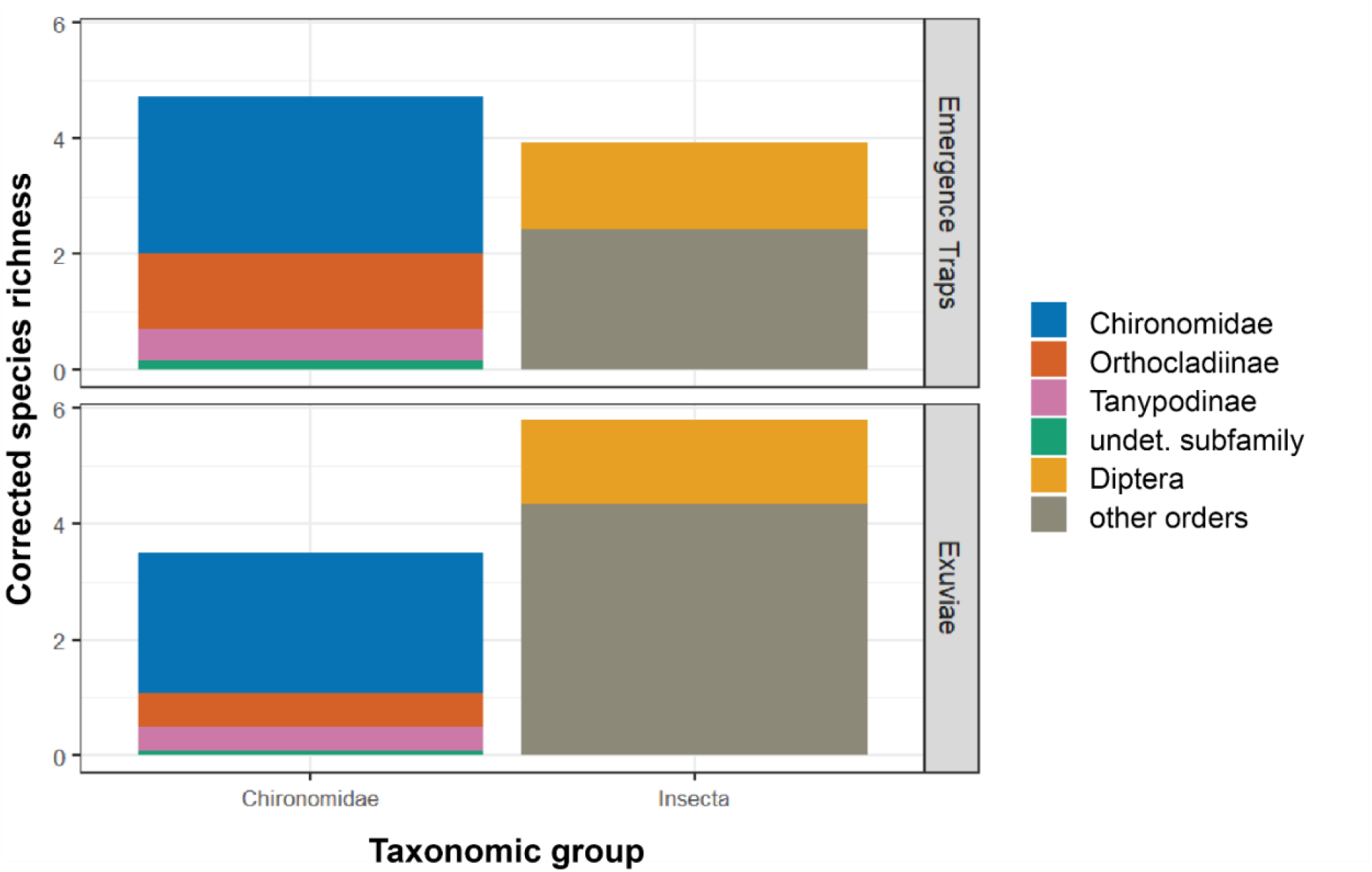
Bar plot of species richness for Chironomidae subfamilies in comparison to all other Insecta and Diptera. Species richness is corrected for the number of samples in each sample type.

**Figure 4.**
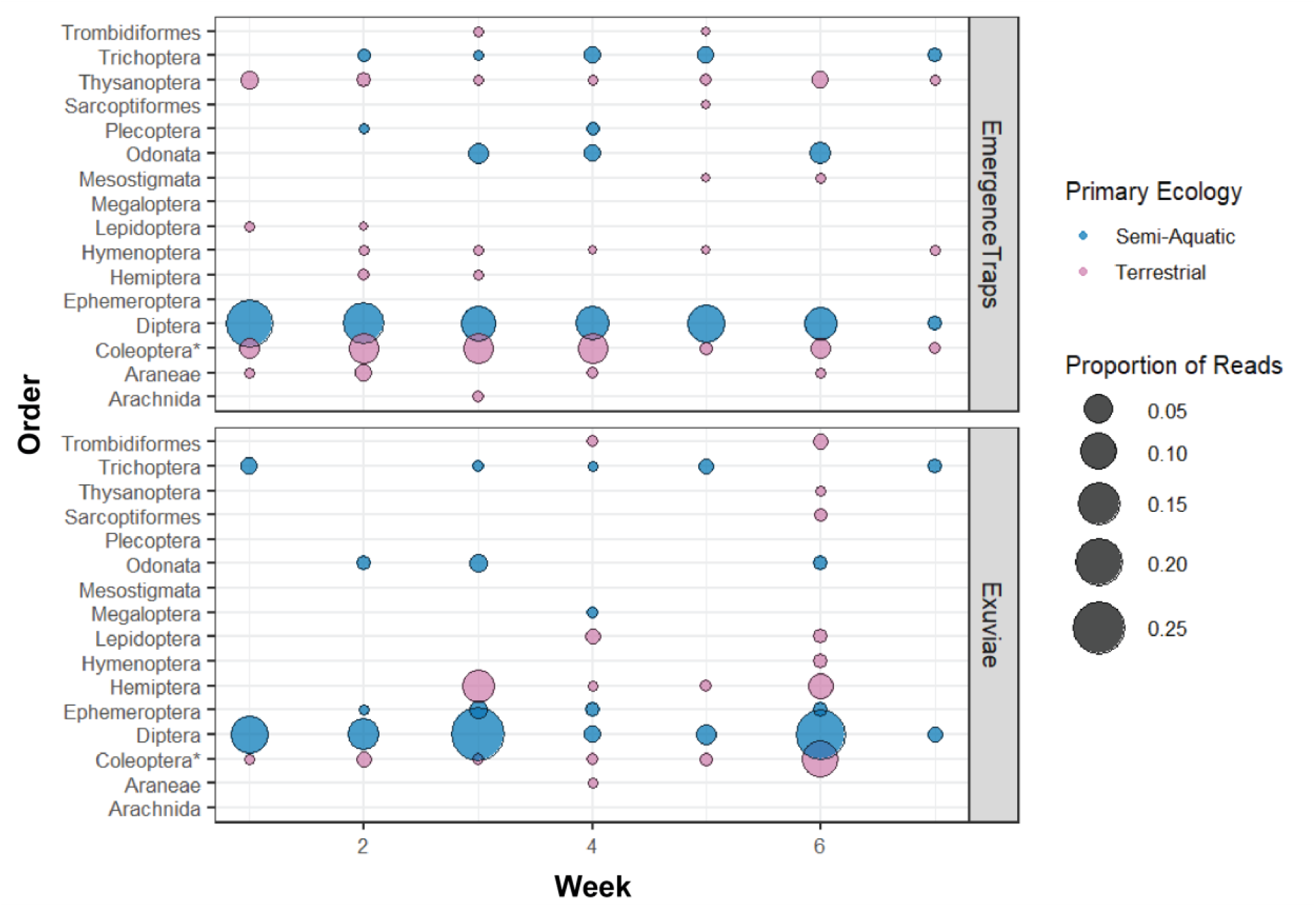
Bubble chart depicting the proportion of reads per sampling week and sample type. The proportional reads are sorted based on insect order and bubbles are sorted and coloured based on the primary ecology of each order, being either semi-aquatic or terrestrial. The asterisk indicates that the order Coleoptera includes a complex mixture of aquatic or terrestrial lifestyles even though grouped as terrestrial.

Overall, chironomids represent the largest taxonomic group in all samples followed by the order Coleoptera which makes up for 13% and 12% of the detections in both emergence and exuviae samples, respectively. In both sample types, all other orders, represent less than 10% of the total detections. Some of these less prevalent taxa were seen at similar rates in both emergence and exuviae samples, e.g. Trichoptera with up 3% and 4% of the total detections, respectively. Orders such as Odonata were seen at a higher rate in exuviae samples (5% vs 1%). The opposite was seen with Thysanoptera (an order of exclusively terrestrial insects) (Mound & Teulon, 1995) which was found in 5% of the detections in emergence traps and in 1% of the exuviae samples.

Although emergence trap samples contained more semi-aquatic species, we found more orders (6) in the exuviae samples (Ephemeroptera, Odonata, Megaloptera, Trichoptera, Lepidoptera and Diptera) (Figure 4). Three of these were also detected in emergence trap samples (Odonata, Trichoptera and Diptera), whereas Ephemeroptera, Megaloptera, and aquatic Lepidoptera were only detected in exuviae samples. The order Plecoptera was only detected using emergence traps (Figures 2 and 4). Beyond the higher number of semi-aquatic orders detected in the exuviae samples, diversity within orders was also higher. For instance, while emergence traps were only able to detect one family of the order Odonata, exuviae samples provided four. Similarily, for Trichoptera, although detected at similar rates in both sample types, we found only one family, Hydropsychidae, in emergence trap samples, but an additional family, Phryganeidae, in exuviae samples. Additionally, the exuviae samples were able to detect two families and two genera from Ephemeroptera and one family and two genera from Lepidoptera, while only one family and one genus was detected for Plecoptera.

## Discussion

This study demonstrates the feasibility of using DNA remnants in exuviae to study semi-aquatic insects and their emergence through metabarcoding. Both quantity and number of reads was low in comparison with bulk samples obtained by emergence traps. This is not unexpected, as exuviae do not contain any DNA but instead, all isolated genomic material is sourced from residual DNA from ecdysis. In addition, much of this residual DNA is lost before collection, due to external environmental conditions such as sunlight, causing degradation of DNA (Sittenthaler et al., 2023). Nevertheless, this does not appear to indicate lower efficiency in terms of the identification of a broad range of taxa as after correcting for the number of samples, both collecting methods showed similar average species richness.

Taxa observed in both emergence trap and exuviae samples were detected at similar rates. Both sample types were dominated by members of the family Chironomidae (Diptera) as well as the orders Trichoptera and Coleoptera. The high richness and relative read count especially for chironomids comes to no surprise as the group is known to be one of the most prevalent taxa of semi-aquatic emerging insects (Ferrington, 2008).

Overall, exuviae samples contained a higher diversity of true semi-aquatic orders (six vs four for emergence samples). Only three orders were detected in both sample types (Odonata, Trichoptera, and Diptera). Within these orders, diversity of exuviae samples was higher for both Odonata and Trichoptera; and lower for Diptera. The higher diversity in Odonata and Trichoptera points to potential limitations of emergence trap sampling. In general, the traps only cover a small area of a waterbody while skimming the surface allows for more coverage and a lower likelihood of missing a local emergence event of e.g. certain caddisfly species. Exuviae samples were collected across the whole constructed wetland and for a large portion of the creek. Moreover, exuviates were free floating, meaning ecdysis did not necessarily occur at the place of collection. Studies have shown that even small changes in environmental conditions, such as substrate, can lead to differences in the emergence patterns of certain semi-aquatic insects (Compson et al., 2013; Gathmann & Williams, 2006). Therefore, it is possible that the location of the emergence traps favoured the collection of certain taxa more than others, while the collection of exuviae covered a larger portion of the body of water, thereby leading to a more general pattern of insect emergence. Another limiting factor might be emergence patterns reported for certain species, e.g. within the Odonata. Previous studies have demonstrated the tendency of some dragonflies to emerge in areas of high vegetation, especially regions of vegetation with higher strata (Hadjoudj et al., 2014). They tend to climb out of the water, onto vegetation, in search for areas exposed to more sunlight, to optimize conditions for the final ecdysis. Emergence traps are not able to accommodate for this behavioural adaption, which could explain the observed low diversity.

Chironomids typically emerge on the surface of the water, and therefore are not prevented from emerging into emergence traps (Cheney et al., 2019) which supports the observed higher diversity in this sampling approach. The collection of exuviae missed some of the diversity present which could be the result of smaller relative body size (Arpellino et al., 2022). Exuviae might have escaped the sieve or the collector’s attention. A potential solution to this might be the use of a large net with fine mesh. That being said, both emergence and exoskeleton samples detected members of the most common subfamilies (Chironominae, Orthocladiinae, and Tanypodinae) showing that despite some limitations metabarcoding of exuviae is capable of detecting a variety of small sized insects.

The larger surface area covered through exuviae collection allowed for the detection of several other orders that contain semi-aquatic species (Megaloptera, Ephemeroptera, and Lepidoptera). Given the comparably small number of semi-aquatic species in Lepidoptera (Pabis, 2018) it is remarkable that several species were found in our surveys. Stone flies (Plecoptera) were only found in very low numbers in emergence traps and not as exuviae. Collection times might have not overlapped with peak emergence time of local species (Singh et al. 2008).

Both methods also provided detections of terrestrial species, e.g. members of the order Thysanoptera, which were found in emergence trap samples. Other groups include various mites and spiders which were also observed in exuviae samples. It can be assumed that these are accidental occurrences. Most of these species are small and could have been passively transported (through wind) to trap structures or onto the surface of the pond or creek.

## Conclusion

Residual DNA found in semi-aquatic insect exuviae can be isolated and used to identify a broad range of taxa by using DNA metabarcoding, making skimming of the water surface and surrounding vegetation a viable method to replace the trap-based capture of insects as they emerge. Such an alternative can also help to reduce the impact of mass collections on semi-aquatic insect populations, which potentially could cause disruptions to vulnerable species. As potentially important new method in the tool box of aquatic entomologists it should be further explored to improve sample size across larger water bodies, and to determine its reliability in different water conditions with varying water chemistry.

## Acknowledgement

This study was supported by funding through the Canada First Research Excellence Fund. It represents a contribution to the University of Guelph Food From Thought research program. We thank the Optimist Club of Kitchener-Waterloo for access to their site at Camp Heidelberg and Al Woodhouse for his help at the site. We are grateful to peer reviewers and editors who provided comments that improved the manuscript.

